# A Single Chimeric Spike Antigen Induces Pan-Sarbecovirus Immunity

**DOI:** 10.1101/2024.11.06.622391

**Authors:** Claudio Counoupas, Paco Pino, Joshua Armitano, Matt D Johansen, Lachlan J Smith, Elizabeth Chan, Caroline Ashley, Eva Estapé, Jean Troyon, Sibel Alca, Stefan Miem-czyk, Nicole G. Hansbro, Scandurra Gabriella, Warwick J. Britton, Thomas Courant, Patrice M. Dubois, Nicolas Collin, V Krishna Mohan, Philip M Hansbro, Maria J Wurm, Florian M. Wurm, Megan Steain, James A. Triccas

**Author notes:** Correspondance: James Triccas.

## Abstract

Next-generation vaccines are required to address the evolving nature of SARS-CoV-2 and to protect against emerging pandemic threats from other coronaviruses. These vaccines should aim to elicit broad-protection, provide long-lasting immunity and facilitate equitable access for all populations. In this study, a panel of chimeric, full-length spike antigens were developed that incorporate mutations from previous, circulating and predicted SARS-CoV-2 variants. The lead candidate (CoVEXS5) was obtained from a high-yield production process in stable CHO cells with purity of >95%, long-term stability and elicitation of broadly cross-reactive neutralising antibodies when delivered to mice in a squalene emulsion adjuvant (Sepivac SWE™). In both mice and hamsters, CoVEXS5 immunisation reduced clinical disease signs, lung inflammation and organ viral titres after SARS-CoV-2 infection, including challenge with the highly immunoevasive Omicron XBB.1.5 subvariant. In mice previously primed with a licenced protein vaccine (NVX-CoV2373), CoVEXS5 could boost T cell immunity, as well as neutralising antibodies levels against viruses from three sarbecoviruses clades. The breadth of sarbecovirus cross-reactivity elicited by CoVEXS5 exceeded that observed after boosting with the NVX-CoV2373 vaccine. These findings highlight the potential of a chimeric spike antigen, formulated in an open-access adjuvant, as a next-generation vaccine candidate to enhance cross-protection against emerging sarbecoviruses in vaccinated populations globally.

## Introduction

The Severe Acute Respiratory Syndrome Coronavirus 2 (SARS-CoV-2) pandemic has underscored the critical role of vaccines in controlling infectious diseases. Since the pandemic’s onset, multiple COVID-19 vaccines have been developed and deployed worldwide^1^. These vaccines, employing diverse technological platforms such as mRNA, viral vectors, protein in adjuvant and inactivated viruses, have significantly reduced infection rates, severe disease and mortality^2^. However the efficacy of licensed vaccines has been constantly challenged by the emergence of SARS-CoV-2 variants, which can evade immune responses and outpace the introduction of updated vaccine formulations^3^. Moreover, the immunity conferred by these vaccines’ wanes over time, which is most apparent with mRNA vaccines, and necessitates multiple booster shots per year for those most at risk^4^. These challenges are compounded by the high costs and logistical hurdles associated with vaccine distribution, particularly in low- and middle-income countries (LMICs), where resources are limited. Thus, there is a need for vaccines that not only offer broad and lasting immunity but are also affordable and easy to distribute globally^5^.

SARS-CoV-2 belongs to the *Sarbecovirus* subgenus of the genus *Betacoronavirus*, which also includes SARS-CoV-1, the cause of the 2002-2004 SARS outbreak. The COVID-19 pandemic intensified surveillance efforts to identify novel coronaviruses present in wildlife around the globe with the capacity to infect humans. In southeast Asia, many coronaviruses closely related to SARS-CoV-1 and -2 have been identified in bats and in other mammals such as pangolins, which may serve as reservoir or bridging hosts; some of these viruses mediate human ACE2-dependent entry and replication in human cells^6,7^. Such viruses pose a threat to human health in the event of a spillover, and the frequency of pathogen emergence from animal reservoirs provides strong evidence that future pandemics similar to COVID-19 will occur^8^. To protect against these pandemic threats, a shift beyond the current reactionary vaccine development paradigm towards preparedness is required. Currently we lack effective tools to control another outbreak of a novel sarbecovirus.

Given its pivotal role in viral entry, the spike protein is the major antigenic component of most commercially available COVID-19 vaccines^9,10^. We have shown that levels of anti-spike neutralising antibodies (nAbs) strongly correlate with protection against symptomatic and severe COVID-19^11,12^. However, evolution of SARS-CoV-2 quickly resulted in variants with increased transmissibility and reduced nAb recognition and consequently vaccine efficacy^13^. Further, current COVID-19 vaccines or prior SARS-CoV-2 infection are unable to effectively neutralise some novel sarbecovirus found in nature, suggesting these viruses pose a risk of zoonotic spillover and adaption to humans^14,15^. In response to these challenges, research is now directed at developing vaccines that are broadly protective, either covering a cross-section of SARS-CoV-2 variants or spanning sarbecovirus clades^16^. Many of these approaches have combined immunogenic regions of the spike antigen, commonly the receptor binding domain (RBD), due to the ease of manufacturing compared to the full-length protein^16^. However full-length spike protein contains a broader selection of immunogenic epitopes; many dominant T cells epitopes fall outside the RBD, and T cell reactivity remains conserved across variants^17^. This may facilitate disease control in the absence of a strong neutralising antibody response.

We have previously demonstrated that full-length trimeric spike antigen produced in an optimised, scalable and chemically defined production process^18^, maintain long-term stability and are highly immunogenic in animal models^19^. We built on this technology to develop a panel of chimeric spike antigens incorporating mutations precited to impact immunogenicity and stability. The lead candidate, CoVEXS5, showed high protein yield, purity, stability and elicited broadly reactive neutralising antibodies in mice and hamsters, reducing clinical disease signs and viral titres after SARS-CoV-2 infection. CoVEXS5 also boosted T cell immunity and sarbecovirus neutralising antibodies in mice previously primed with an approved COVID-19 vaccine, highlighting its potential as a next-generation vaccine for broad sarbecovirus protection.

## Results

### Selection of broadly immunogenic Chimeric Spike Antigens

We sought to identify chimeric spike antigens that are highly immunogenic and produce cross reactive immunity to diverse SARS-CoV-2 variants. Forty chimeric spike constructs were initially designed, which was downselected to 13 constructs after modelling for trimer formation (Figure 1A). All protein antigens are chimeric spike ectodomains and include 6 proline mutations to improve expression and thermostability ^9^ and lacked the furin cleavage site. Additional spike mutations were selected based on mutations known or predicted to impact infectivity and/or immunogenicity^20^, or identified through *in vitro* evolution studies^21^ (Figure 1C). Plasmids encoding each of the 13 constructs were transfected into CHOEXpress® cells to generate stable pools. Twelve of 13 (92%) constructs resulted in protein expression. Seven of these chimeric spike constructs were expressed as trimeric molecules and had a range of initial productivity, from 50 mg/L to 400 mg/L, which increased upon subsequent optimisation (Figure 1B). The 7 detectable constructs were purified (one-step affinity resin) and all proteins produced displayed >90% purity, as measured by HPLC-SEC. CSA05 and CSA07 were produced at the highest level (Figure 1B).

**Figure 1.**
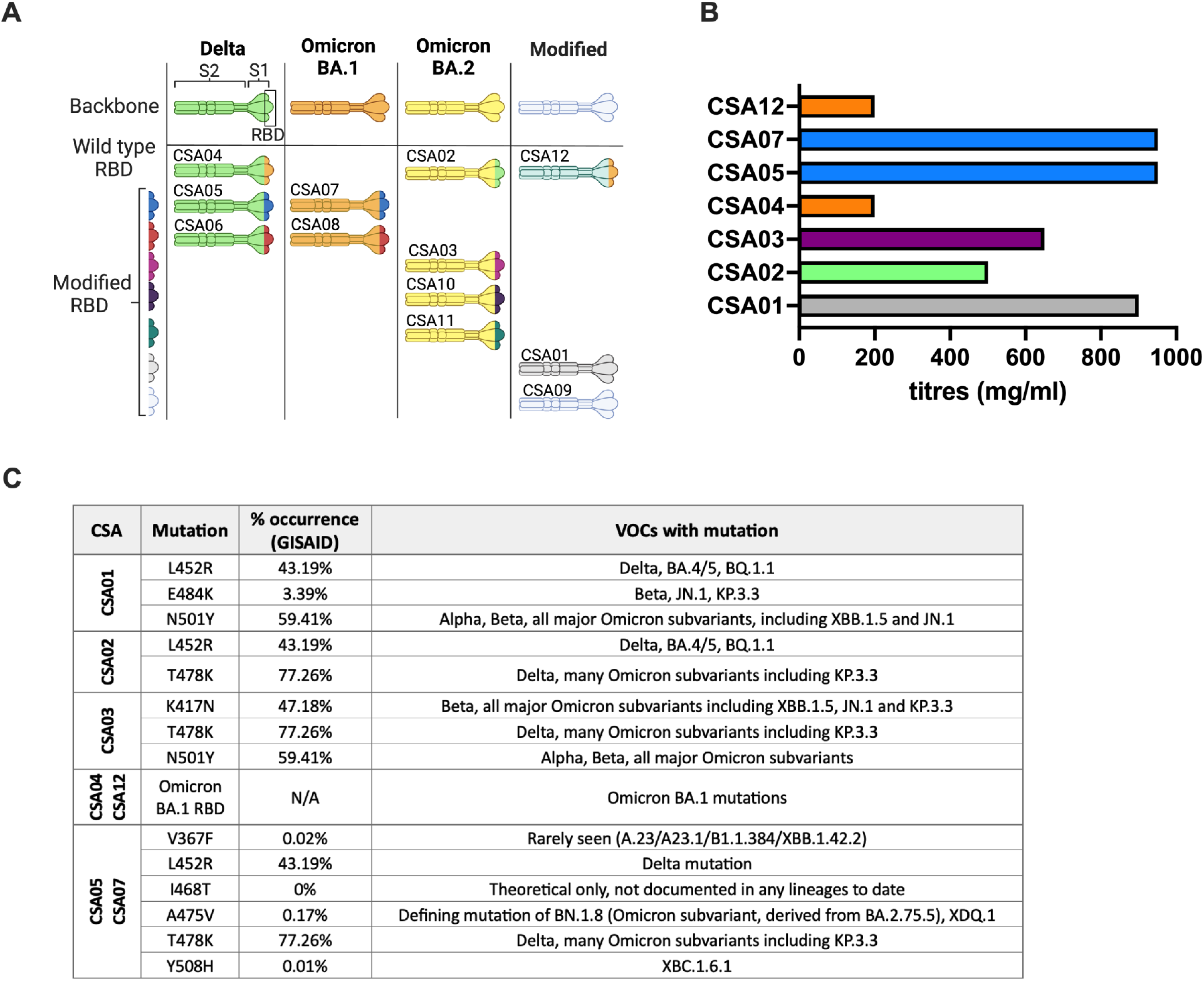
Development of chimeric spike antigens. **A**. Schematic representations of the chimeric spike proteins (CSAs), colour coded to show their relationship to VOC or between themselves. Colours are chosen arbitrarily but identical colour reflects identical sequences. **B**. Manufacturability of various of CSAs in CHO cells: titers reached after 14 days small-scale fed-batch evaluation (CSAs not represented e.g. CSA09, could not be expressed to a usable titer). **C**. Mutations present in CSAs.

The 7 chimeric spike antigens (CSA) were formulated in Sepivac SWE™, an open access oil-in-water emulsion and delivered to mice (intramuscularly (i.m.) at days 1 and 21). At 42 days post the first immunisation, nAb titres (IU/ml) in plasma were determined using a panel of spike-pseudotyped viruses. No nAbs were observed in antigen or adjuvant alone groups (not shown). As represented by the heatmap of nAb responses (Figure 2A), most CSAs demonstrated appreciable nAb levels against the majority of variants tested, although CSA04 and CSA12 displayed limited induction of cross-neutralising nAbs, and both these spike variantsshare the Omicron BA.1 RBD, which may account for their poor cross-reactivity. Notably, CSA05 and CSA07 displayed strong activity against early pandemic variants (Ancestral, Alpha, Beta, Delta) and all Omicron subvariants tested (Figure 2B). Conversely, vaccination with Ancestral spike only showed good nAb activity against early pandemic variants (non-Omicron), while BA.1 spike immunisation resulted in restricted neutralisation of Omicron subvariants (Figure 2B). To select the CSA candidates for progression, we used a ranking system that examines the area under the curve (AUC) when nAb levels are plotted against the genetic distance between SARS-CoV-2 variants^22^. This allows us to devise a score for each candidate based on their ability to neutralise a wide cross-section of variants (Figure 2B with individual plots shown in Figure 2C). The top candidate based on this ranking was CSA05, which also showed good manufacturability (Figure 1B) and long-term stability (Supplementary Table 1). CSA05 combined with the Sepivac SWE^™^ adjuvant was termed CoVEXS5 and progressed to further analysis.

**Figure 2.**
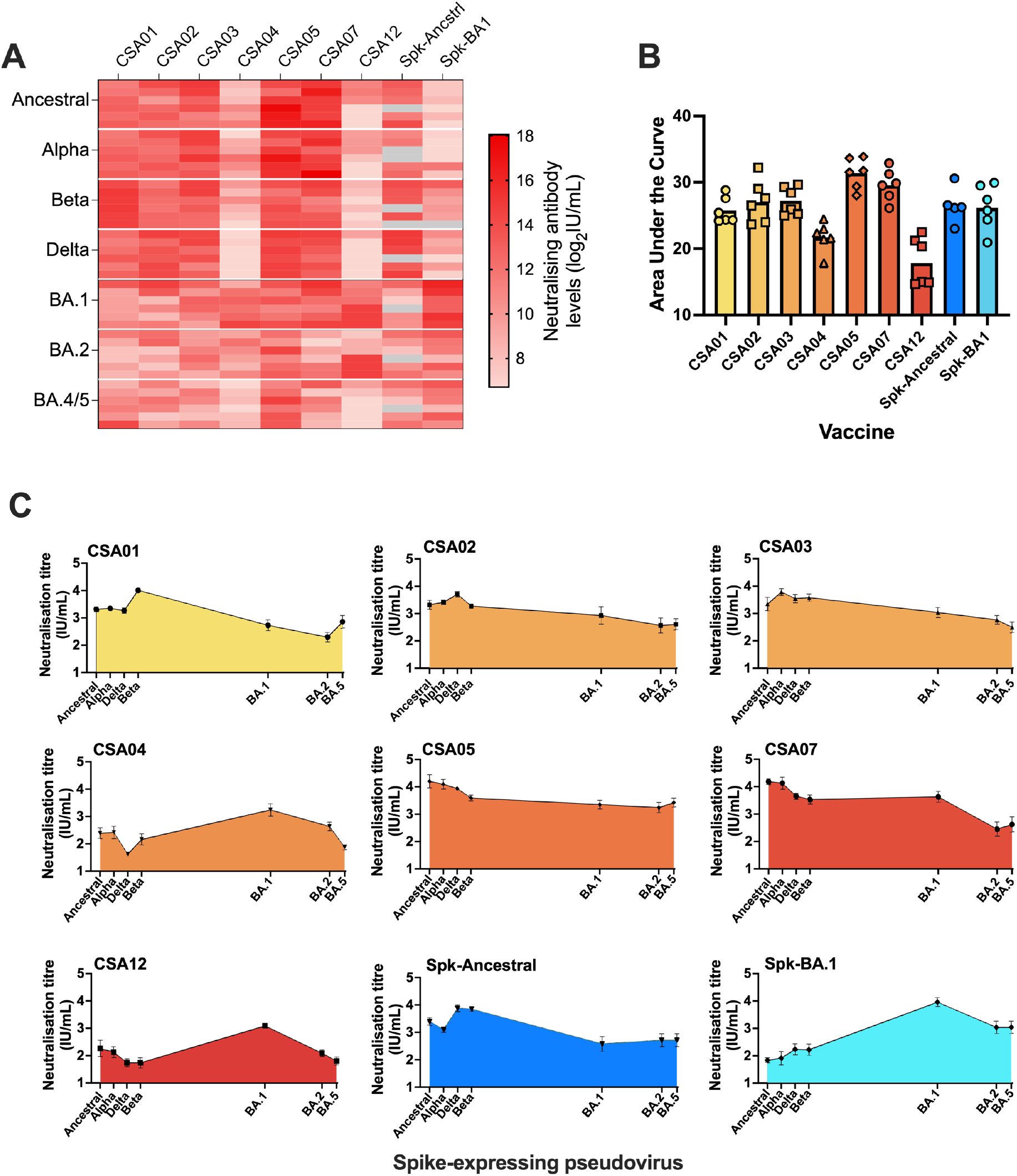
Selection of chimeric spike antigens with broad neutralising activity. **A**. Neutralisation of pseudoviruses by plasma from mice immunised with CSA candidates. Mice were vaccinated twice, 3 weeks apart with 5 *μ*g of each spike antigen formulated in SWE adjuvant. Twenty-one days after the last dose, nAb titres against pseudotyped viruses was determined. **B**. Area under the curve (AUC) to demonstrate breadth of nAbs across different variants. Phylogenetic difference is based on variant RBD amino acid sequence. The results are expressed as the number of amino acid substitutions per 100 amino acids, relative to ancestral virus. **C**. Individual plots for AUC analysis.

### Protection against virulent SARS-CoV-2 in mice and hamsters

To determinedetermine if CoVEXS5-induced immunity was protective against SARS-CoV-2 infection, K18-hACE2 were vaccinated twice, 3 weeks apart with CoVEXS5, at 3 concentrations of CSA05 formulated 1:1 vol/vol with Sepivac SWE^™^. Twenty-eight days later mice were challenged intranasally with SARS-CoV-2 (Delta variant; Figure 3A). Mice sham-vaccinated with PBS succumbed to infection within 6 days displaying substantial deterioration in their condition (Figure 3B) with ∼15% weight loss (Figure 3C). These events led to extensive lung inflammation with substantial increases in inflammatory cells in the airways (Figure 3D). Vaccination with all doses of CoVEXS5 completely protected against infection, with no observable change in clinical score (Figure 3B) or weight loss (Figure 3C). Vaccination with higher doses of CoVEXS5 reduced BALF inflammation, with partial reduction at the lower dose of 0.1 *μ*g (Figure 3D). However, all CoVEXS5 doses resulted in no detectable infectious virus in the airways, lungs and brain (Figure 2E). Thus, CoVEXS5 is sufficient to completely protect mice from the development of COVID-19 disease manifestations, to neutralise infectious SARS-CoV-2, and prevent pathogenic inflammation in the lung.

**Figure 3.**
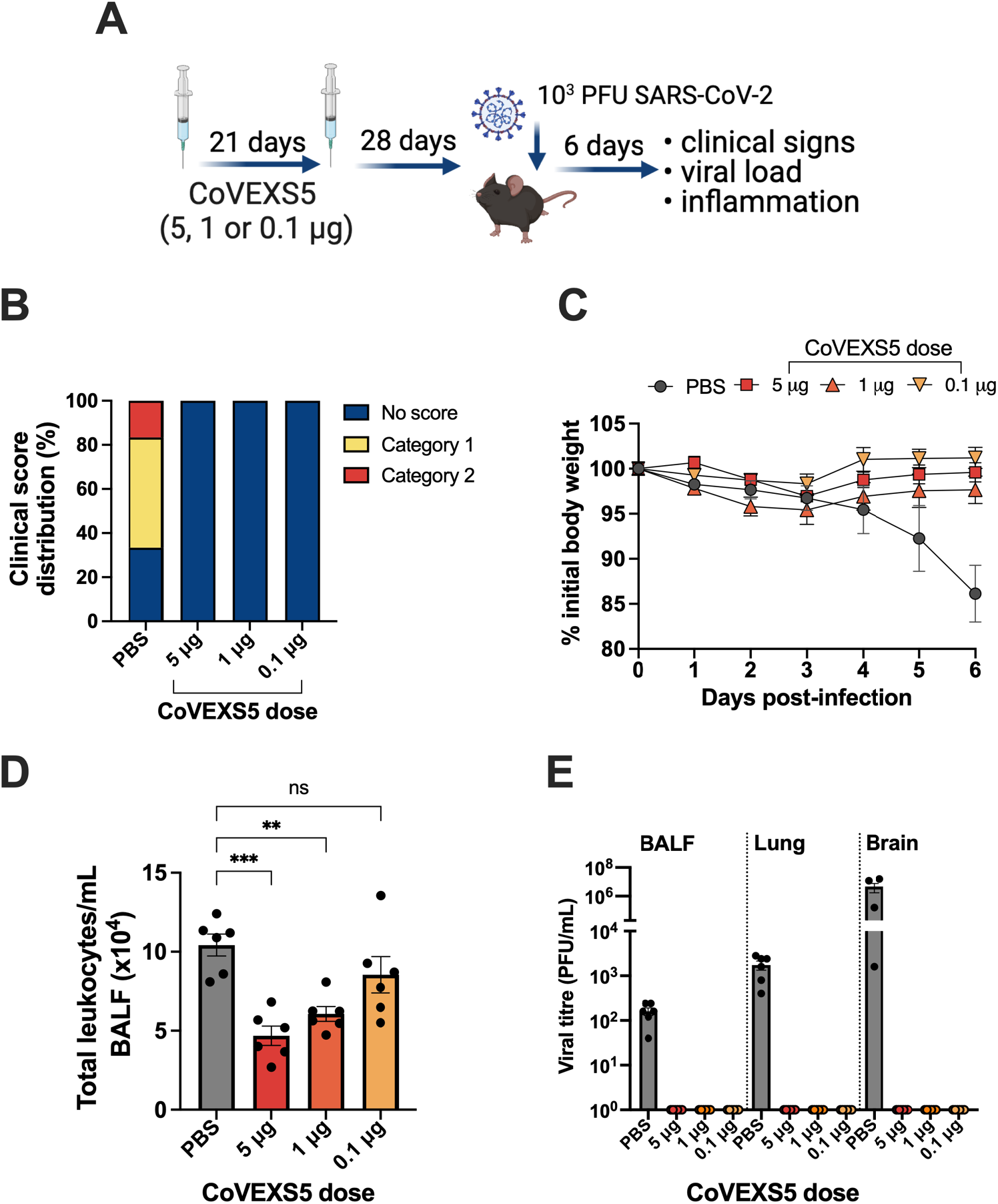
CoVEXS5 protects against severe SARS-CoV-2 infection in mice. **A**. K18-hACE2 mice (n=6) were immunised twice with sham (PBS) or CoVEXS5 at 5, 1 or 0.1 *μ*g of CSA05 formulated 1:1 with SWE and at 28 days challenged with 10^3^ PFU SARS-CoV-2 Delta. **B**. Clinical scores at day 6 post-infection **C**. Percentage of initial body weight loss in male K18-hACE2 mice (n=6/group). **D**. Total inflammatory cells in bronchoalveolar lavage fluid (BALF). **E**. Viral titres in BALF, lung or brain homogenates were determined using plaque assay. Significant differences between placebo and vaccinated group were determined by one-way ANOVA with Šidák’s multiple comparison test; *p<0.05, **p<0.01.

To determine if efficacy extended across species and to more diverse SARS-CoV-2 variants, hamsters were immunised with two doses of CoVEXS5 or Novavax NVX-CoV2373 as a comparator (Figure 4A). Both vaccines significantly protected animals from weight loss after challenge with the Delta virus (Figure 4B) or the highly immune-evasive Omicron XBB.1.5 virus (Figure 4C), compared to placebo controls. In vaccinated animals challenged with the Delta variant, there was complete clearance of virus in the majority of animals in both nasal turbinates and the lung, irrespective of the CoVEXS5 dose used (Figure 4D). In animals challenged with Omicron XBB.1.5, all vaccines significantly reduced viral load compared to sham-vaccinated animals, although to a lesser extent than the protection observed with Delta variant challenge. A greater reduction in viral load was seen with the 20 *μ*g compared to the 5 *μ*g dose of CoVEXS5. Thus, CoVEXS5 afforded protection against two divergent SARS-CoV-2 variants in hamsters.

**Figure 4.**
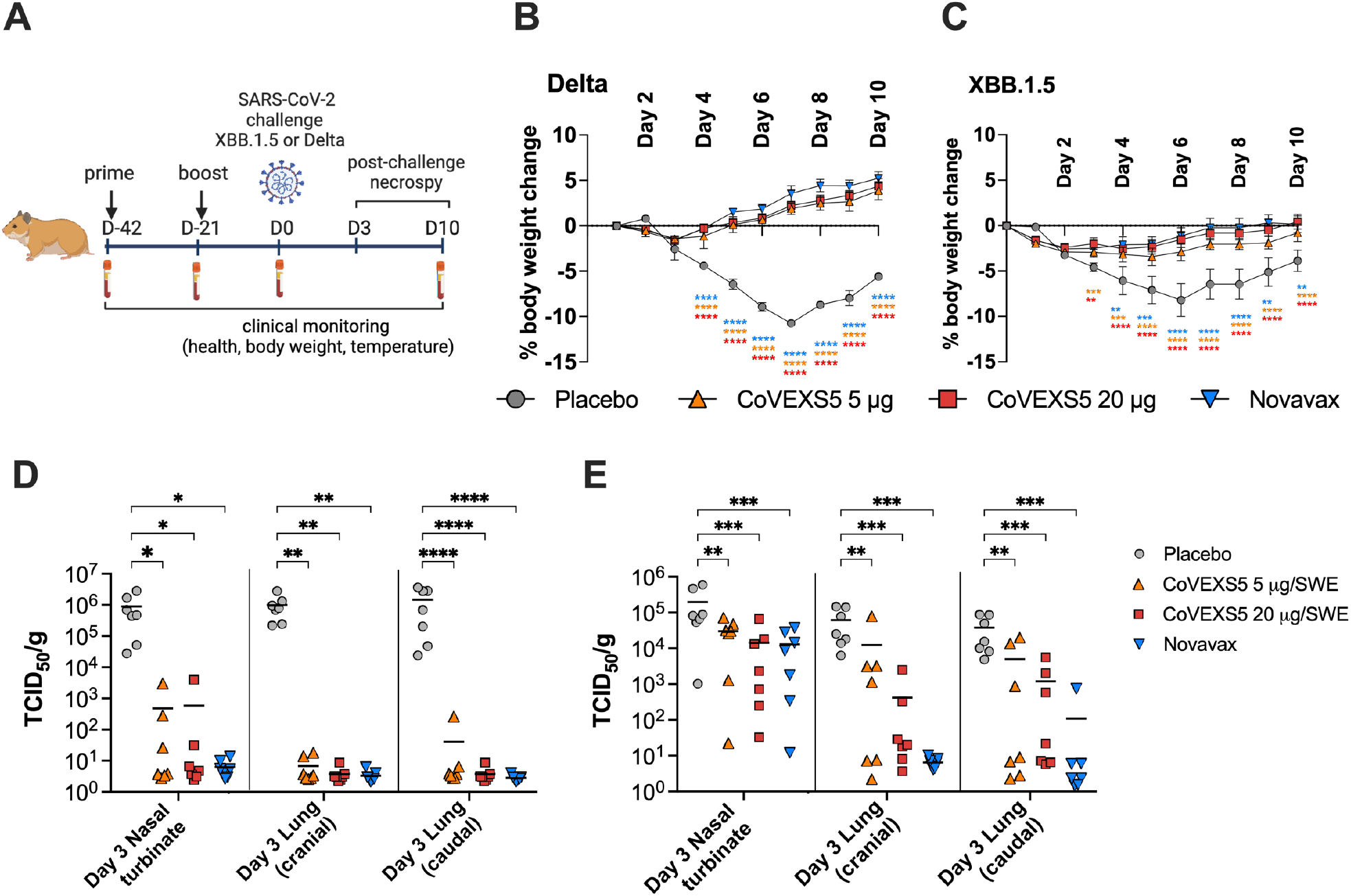
Protection of CoVEXS5-vaccinatedvaccinated hamsters against infection with divergent SARS-CoV-2 variants. Hamsters were vaccinated with CoVEXS5 (5 *μ*g or 20 *μ*g) or NVX-CoV2373 (5 *μ*g) as shown in Panel **A**, and infected with either SARS-CoV-2 Delta or XBB.1.5 at 21 days post-final vaccination. Change in body weight after challenge is shown for Delta (**B**) or XBB.1.5 (**C**) challenge. Also shown are viral loads in nasal turbinates and the lung at Day 3 post challenge (Delta, **D**; XBB.1.5, **E**). Significant differences between placebo and vaccinated group were determined by two-way ANOVA with Dunnett’s multiple comparison test; *p<0.05, **p<0.01, ***p<0.001, ****p<0.0001.

### CoVEXS5 boosts pre-existing immunity to provide broad sarbecovirus immunity

Given the majority of the population has now been vaccinated and/or infected with SARS-CoV-2, we determined if the lead candidate, CoVEXS5, could boost pre-existing immunity. Mice were primed with an approved COVID-19 vaccine (NVX-CoV2373 or NVX) and left for an 18-week period to allow responses to wane (Figure 5A). NAbs from vaccinated mice were assessed against a representative panel of 6 spike-expressing pseudoviruses. After peaking at 4-6 weeks post-vaccination (1-3 week post the 2^nd^ dose), nAbs declined against all pseudoviruses examined (Figure 5B).

**Figure 5.**
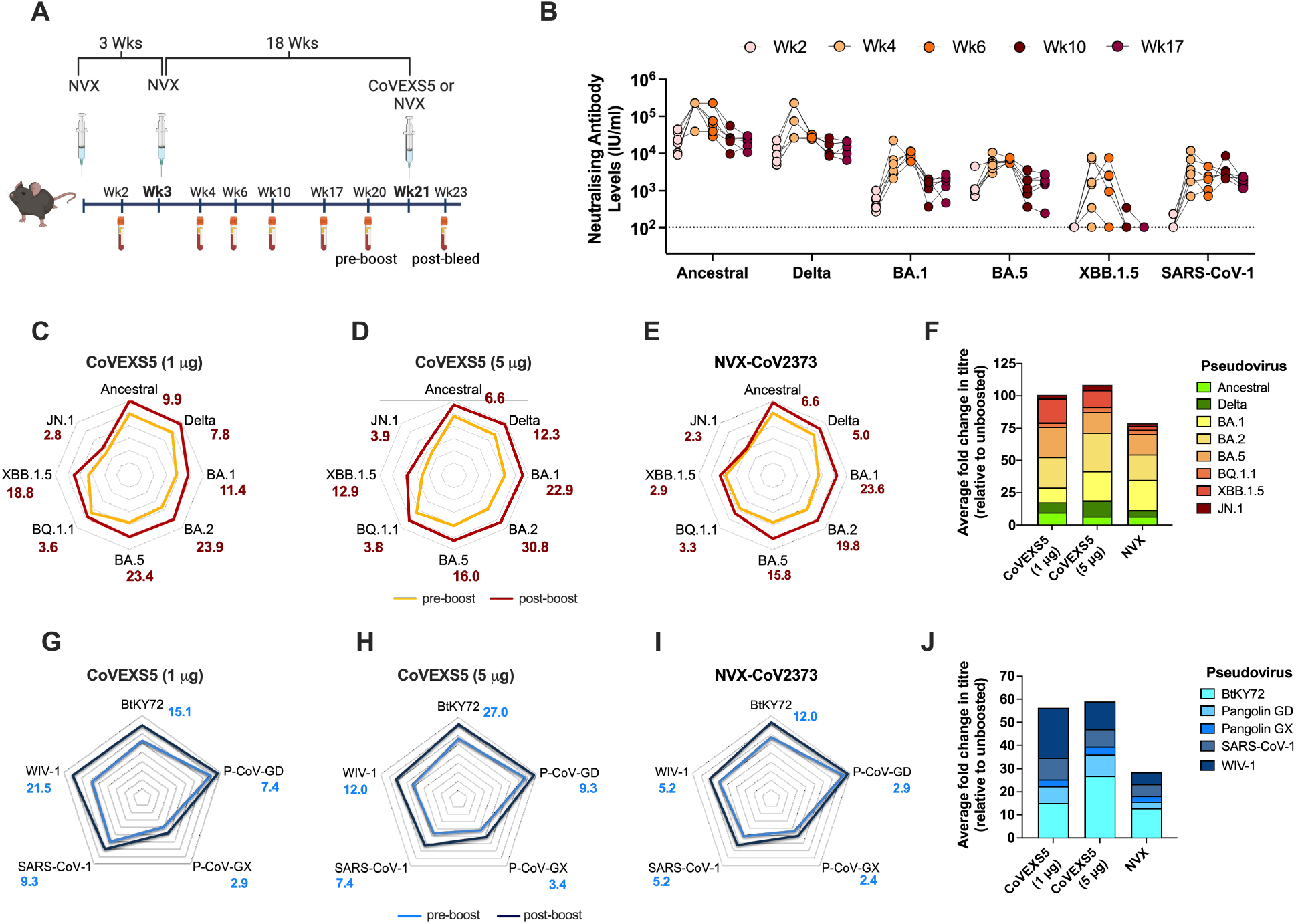
Broad sarbecovirus immunity after CoVEXS5-boosting in mice. A. C57BL/6 mice (n=6) were vaccinated i.m. with 2 doses of NVX-CoV2373 (NVX) at 3 weeks apart (1 *μ*g per dose) and then boosted 18 weeks later with 1 or 5 *μ*g of CoVEXS5, or 1 *μ*g of NVX. B. Pre-boost kinetics after NVX mmunisation for a selection of spike-pseudotyped viruses. Pre-and post-nAb levels were determined using a panel of spike-pseudotyped viruses representing early SARS-CoV-2 viruses (C-E) or non-SARS-CoV-2 sarebecoviruses (G-I) in mice vaccinated with CoVEXS5 1 *μ*g, (C,G), CoVEXS5 5 *μ*g (D,H) or NVX (E, I). Numbers represent the fold change between pre- and post-boost nAb levels. The stack bar graphs show the fold change for each SARS-CoV-2 variant (F) or non-SARS-CoV-2 sarbecoviruses (J). For C-E and G-I, individual data points are shown in Supplementary Table 2.

To define the impact of boosting with CoVEXS5, NVX-vaccinated mice were boosted with SWE-adjuvanted vaccine containing either 1 *μ*g or 5 *μ*g of CSA05, or boosted with an additional dose of NVX (Figure 5A). Boosting with CoVEXS5 increased nAb levels against early pandemic variants (Ancestral, Delta) and Omicron subvariants, irrespective of the dose of vaccine used (Figure 5C-E). This boosting effect ranged from approximately 3-fold to 30-fold, dependent on the variant examined. Encouragingly, the impact of boosting with CoVEXS5 was apparent for the highly immune-evasive BQ.1.1, XBB.1.5 and JN.1 subvariants (Figure 5C-E). NAb responses against a panel of five non-SARS-CoV-2 sarbecoviruses from Clade 1A (SARS-CoV-1, WIV-1), Clade 1b (Pangolin-GD, Pangolin-GX) and Clade 3 (BtKY72) were also assessed. These viruses were selected as they presented a cross-section of sarbecoviruses, and their spike sequences were not used for the design of the CSAs described in Figure 1, to determine the broad utility of the vaccine. CoVEXS5 immunisation resulted in increased nAbs against all viruses, notably BtKY72 (27-fold) and WIV-1 (12-fold), with the 5 *μ*g dose eliciting the highest levels of cross-reactive nAbs (5G-J). The cumulative level of nAb boosting was greater for the CoVEXS5 vaccines compared to NVX, for both SARS-CoV-2 variants (Figure 5F) and non-SARS-CoV-2 sarbecoviruses (Figure 5J).

T cells responses after boosting were also examined, by determining the level of cytokine release by CD4^+^ or CD8^+^ T cells from lung or spleen cells after stimulation with spike protein. CoVEXS5 at both doses was able to boost T cell responses after NVX prime, which was most apparent for CD4^+^ T cells secreting IL-2 or IL-5 in the lung (Figure 6A, 6B) or spleen (Figure 6E, F). The proportion of CD8 T cells secreting IFN-*γ* was significantly increased compared to naïve mice in the lung (Figure 6C, F) and spleen (Figure 6E, F) for all vaccines, with the low dose of CoVEXS5 resulting in the greatest proportion of CD8 T cells secreting TNF at both sites, compared to unvaccinated animals. Thus, CSA05 is able to stimulate both humoral and T cell responses following priming with approved COVID-19 vaccines, resulting in augmented and broad-spectrum immunity against diverse SARS-CoV-2 variants and other sarbecoviruses.

**Figure 6.**
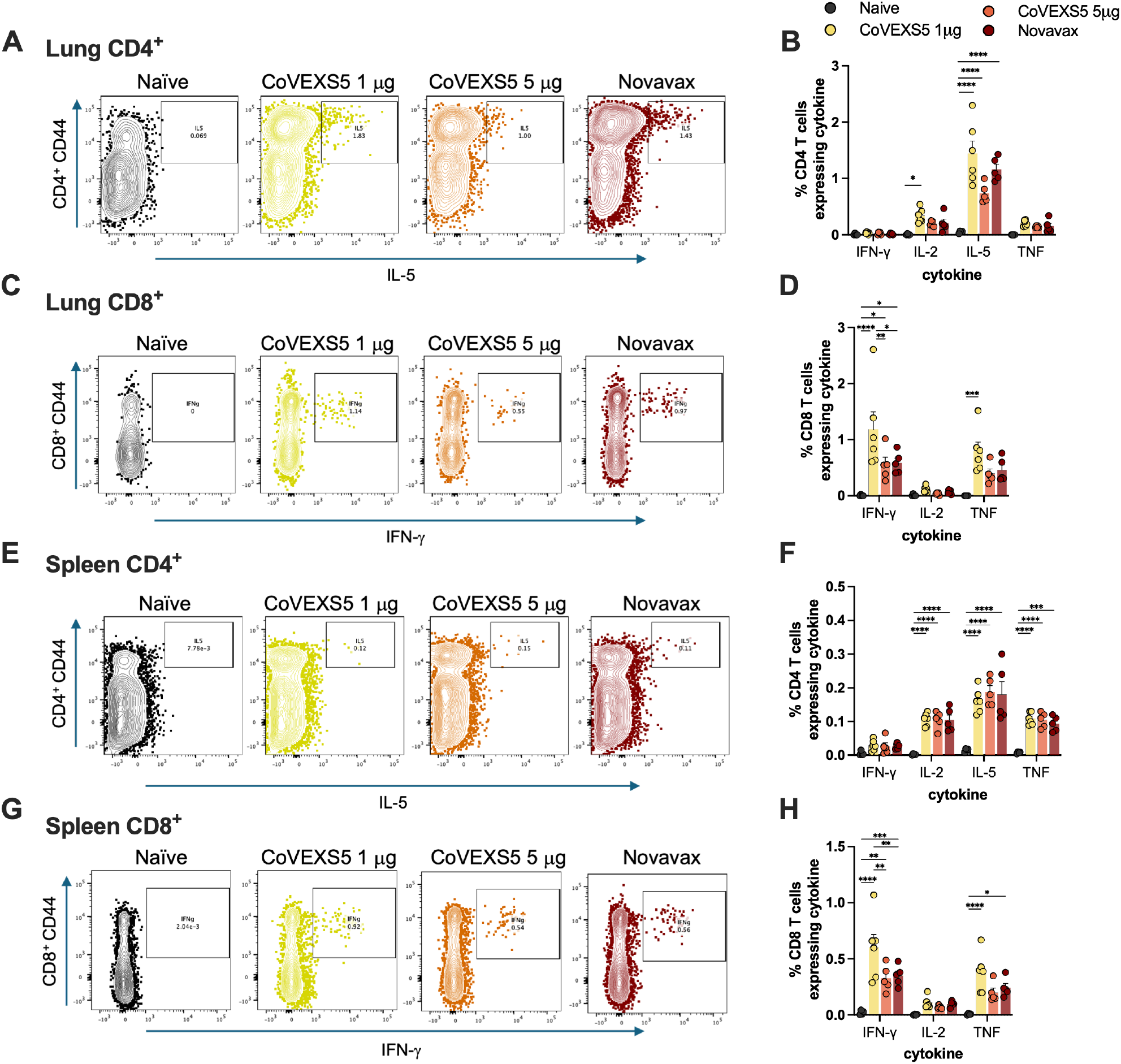
Boosting of T cell responses with CoVEXS5. Mice were vaccinated as in Figure 5. At 14 days post boosting, lung (A, B) or spleen (C, D) cells were restimulated with 5 *μ*g/mL of SARS-CoV-2 ancestral spike protein and the frequency of CD4^+^ (A, C) or CD8^+^ (B, D) T cells expressing cytokines was determined by flow cytometry. Significant differences between placebo and vaccinated group were determined by two-way ANOVA with Tukey’s multiple comparison test; *p<0.05, **p<0.01, ***p<0.001, ****p<0.0001.

## Discussion

The ongoing evolution of SARS-CoV-2 results in the emergence of new viral variants capable of evading pre-existing immunity, necessitating the continual updating of vaccines. A vast pool of antigenically distinct sarbecoviruses also circulates in nature, posing a further risk of zoonotic spillovers to humans. Thus, the development of next-generation vaccines that can elicit broad sarbecovirus protection and can be easily distributed is essential.

The vaccine candidate described in this report, CoVEXS5, combines a novel highly stable chimeric spike antigen with the open-access adjuvant Sepivac SWE^™^. The spike antigen in CoVEXS5 has been engineered not only for stability and optimal expression but also to incorporate key mutations from other SARS-CoV-2 variants, enhancing the breadth of protection against evolving strains. Importantly, CoVEXS5 elicits robust and cross-reactive immune responses against a range of SARS-CoV-2 variants, including the highly immune-evasive Omicron XBB.1.5 and JN.1 subvariants, and against a broad range of diverse sarbecoviruses. The protective efficacy of CoVEXS5 was demonstrated in both mice (Figure 3) and hamsters (Figure 4) against diverse SARS-CoV-2 variants. These findings are promising for the potential use of CoVEXS5 in humans, where cross-species efficacy is a valuable indicator of broad protective capabilities.

Approved COVID-19 vaccines are based on single or bivalent spike antigens and are unable to maintain efficacy over time and as new variants have emerged^13^. In addition, they require updated formulations and frequent booster doses to sustain protection [2, 3]. We combined mutations into fulllength spike proteins to identify stable and cross-reactive antigens (Figure 1). Some chimeric spikes contained variant-based RBD sequences, such as CSA04 and CSA12, which had a BA.1-based RBD. However, both CSA04 and CSA12 performed poorly against non-Omicron variants; in fact, these chimeric spikes ranked lower than the wild-type BA.1 spike antigen, showing that certain combination of mutations are detrimental to immunogenicity (Figure 2). The two best ranked chimeric spikes, CSA05 and CSA07, contained identical RBD mutations, derived from various circulating variants, or mutations that at the time of antigen design (around the time of BA.1/BA.2 emergence) were predicted to arise. Encouragingly, the RBDs of CSA05 and CSA07 contain a mix of mutations that either have been preserved in the majority of Omicron sublineages (TbK^23^), have independently arisen in later Omicron lineages (L452R in BA.4/5^24^) or that are being increasingly observed in new Omicron sublineages such as JN.1 (A475V^25^). The broad cross-protection seen with CSA05 when formulated into CoVEX5, and superiority in generating nAbs against a range of sarbecoviruses compared to the ancestral spike-based vaccine NVX, validates our chimeric antigen approach.

The CHOEXpress® platform we use allowed high-yield production of chimeric antigens, with the lead CSA05 protein approaching yields of 1 g/L (Figure 1) and demonstrating long-term stability at 4°C (Supplementary Table 1). This is particularly crucial for global vaccination efforts, especially in resource-limited settings where cold chain logistics pose significant challenges. When combined with the open-access Sepivac SWE™ adjuvant, this represents a significant step towards facilitating vaccine accessibility. The SWE adjuvant is a squalene oil-in-water emulsion, similar in composition to MF-59^®^, a class of adjuvants with a strong safety record in humans^26^. Importantly, we observed cross-reactive immunity with a single chimeric antigen. This differs to other approaches to develop pan-sarbecovirus vaccines, that require multiple protein components to achieve broad immune responses^27,28^. Single chimeric antigens will have reduced complexity and cost of vaccine development, making them suitable for broad global distribution.

The majority of people across the globe have been vaccinated with at least a primary series of COVID-19 vaccines encoding ancestral spike antigen. We approximated this scenario in mice by priming with the NVX-CoV2373 vaccine and showed that boosting with CoVEXS5 significantly increased nAb levels against various SARS-CoV-2 variants and antigenically distinct sarbecoviruses from different lineages (Figure 5). Notably, the overall effect of boosting was enhanced compared to boosting with a third dose of NVX-CoV2373, particularly for the non-SARS-CoV-2 sarbecoviruses. This suggest that the modified chimeric spike used here is able to present a broader repertoire of cross-reactive epitopes than ancestral spike antigen. This could relate to the concept of original antigenic sin, where immunity to a first antigen stimulus impacts the induction of *de novo* responses^29^, which can be partly overcome by vaccinating with more antigenically distinct antigens^30^. Alternatively, the use of the Sepivac SWE^™^ adjuvant may also facilitate such cross-reactive responses, an effect seen when this adjuvant was combined with bivalent and trivalent spike antigen combinations^31^. Strong CD4^+^ and CD8^+^ T cell responses were also observed after CoVESX5 delivery (Figure 6), which may serve as an additional line of immune defence to limit disease severity^17^. The ability to enhance both humoral and cellular immunity highlights the potential of CoVEXS5 as a universal booster vaccine, with the potential to provide protection against further zoonotic spillovers of diverse sarbecoviruses.

In conclusion, CoVEXS5 represents a significant step forward in the development of an accessible pan-sarbecovirus vaccine. The robust immune responses elicited by CoVEXS5, its ability to boost preexisting immunity, and its efficacy in diverse animal models underscore its potential as a next-generation vaccine. Future research should focus on clinical trials to confirm these findings in humans and explore the full potential of CoVEXS5 in pandemic preparedness. Additionally, our chimeric antigen design may be applied to other families of viruses to provide cross-protection against antigenically diverse members.

## Methods

### Chimeric Spike design and expression

Sequences of Spike proteins of available variants were collected from public database (GISAID). Selected mutations from variants and predicted potential mutations were incorporated into a Hexapro spike sequence ‘backbone’ and sequences were run through ROBETTA^32^ and SWISS-MODEL^33^ for homology structure prediction for trimer formation. DNA constructs were synthetised by ATUM (Menlo Park, USA) and provided with CMC relevant documentation. The furin cleavage-site RRAR was mutated to be non-functional. The transmembrane domain and the C-terminal intracellular tail were removed and replaced by a T4 foldon sequence in trimer designs^34^. A four amino acid C-terminal ‘C-tag’ was added to purify chimeras that could not bind commercially available resins^35^.

Transfection and culturing of CHOExpress® cells (ExcellGene SA, Monthey) was performed as previously described^18^. After transfections, suspension cells were selected with puromycin, stable pools selected with high secretion and clonal cell lines were obtained by image-assisted cell distribution (f-sight, Cytena GmbH, Freiburg). The lead clonal cell lines were used for scale-up in an optimised fed-batch process at the 0.2 L, 10 L and 50 L bioreactor scale of operation. The production medium utilised was EX-CELL® Advanced™ CHO Fed-batch medium (Merck-Sigma). Bioreactors were seeded at a density of 5×10^5^ cells/mL and maintained at 37°C for 4 days. Production culture fluids were harvested, clarified using Harvest RC (3MTM), and proteins purified via affinity chromatography. The loading, washing, and elution steps were performed on an NGL COVID-19 resin (Repligen, Waltham, USA) as per the manufacturer’s recommendations. The eluted product was further purified using Capto adhere anion exchange resin (Cytiva), followed by tangential flow filtration with a 100 kD cut-off to isolate trimeric spike proteins.

### Mouse Immunisation

Female C57BL/6 mice (6-8 weeks of age) were purchased from Australian BioResources (Moss Vale, Australia); K18-hACE2 mice were bread as hemizygous at Centenary Institute, Newtown, Australia. All mice were housed at the Centenary Institute in specific pathogen-free conditions. All mouse experiments were performed according to ethical guidelines as set out by the Sydney Local Health District (SLHD) Animal Ethics and Welfare Committee (ethics approval number 2020-009).

Mice were immunised twice, three weeks apart intramuscularly (i.m.). Each vaccine contained 5, 1 or 0.1 *μ*g of different stabilised full length spike antigen (chimeric, ancestral or BA1) in endotoxin free PBS with 25 *μ*l of Sepivac SWE™ (SWE) adjuvant (1:1 volume ratio; Seppic, France). For boosting experiments mice received a prime with 100 *μ*l of Novavax NVX-CoV2373 vaccine twice, three weeks apart i.m. (corresponding to 1*μ*g of ancestral spike protein), rested 18 weeks before being boosted with chimeric spike in SWE (1-5 *μ*g of antigen) or Novavax NVX-CoV2373.

### Pseudovirus Neutralisation Assays

Replication-deficient SARS-CoV-2 Spike pseudotyped lentivirus particles were generated by cotransfecting GFP-luciferase, blue or LSSmOrange vector and a spike expression construct together with lentivirus packaging and helper plasmids into 293T cells using Fugene HD (Promega, Wisconsin, USA) as previously described^36^. To determine neutralising antibody titers, pseudovirus particles were incubated with serially diluted plasma samples at 37°C for 1 hr prior to spinoculation (800xg) of ACE2 over-expressing 293T cells. Seventy-two hours post-transduction, cells were fixed and stained with Syto60 (SYTO™ 60 Red Fluorescent Nucleic Acid Stain, Invitrogen) as per the manufacturers instruction’s, imaged used an Opera Phenix high content screening system (Perkin Elmer, Massachusetts, USA) and the percentage of GFP, blue or LSSmOrange positive cells was enumerated (Harmony® high-content analysis software, Perkin Elmer). To determine the breath of neutralisation across variants, neutralisation titres for each were plotted on the y-axis, while the x-axis displayed the genetic distance of the RBD for each pseudotyped spike virus relative to ancestral RBD. This generated a unique curve for each mouse plasma sample. Area under the curve was measured for each curve generated using GraphPad Prism.

### Flow Cytometry

Spleen and lung samples were processed as described previously^37^. To assess spike-specific cytokine induction by T cells, murine lung or spleen cells were stimulated for 4 hrs with Ancestral spike (5 *μ*g/mL) and then supplemented with Protein Transport Inhibitor cocktail (Life Technologies) for a further 10-12 hrs. Cells were surface stained with Fixable Blue Dead Cell Stain (Life Technologies) and marker-specific fluorochrome-labeled antibodies. Cells were then fixed and permeabilised using the BD Cytofix/Cytoperm^™^ kit (Beckton Dickinson, New Jersey, USA) according to the manufacturer’s protocol and intracellular staining was performed to detect cytokines IFN-*γ*, IL-5, IL-2, TNF. All samples were acquired on a BD-LSRII and assessed using FlowJo^™^ analysis software v10.6 (Treestar, Oregon, USA).

### Mouse SARS-Cov-2 challenge

Male hemizygous K18-hACE2 mice were challenged as described previously ^36^. Briefly, Mice were anaesthetised with isoflurane followed by intranasal challenge with 10^3^ PFU SARS-CoV-2 (Delta strain). Mice were weighed and monitored daily. At day 6 post-infection, mice were euthanised and BALF was collected and enumerated using a haemocytometer (Sigma-Aldrich, USA). Tissue was homogenised using a gentle MACS tissue homogeniser, after which homogenates were centrifuged (300 *g*, 7 min) to pellet cells, followed by collection of supernatants for viral quantification by plaque assay. VeroE6 cells (CellBank Australia, Australia) were grown in Dulbecco’s Modified Eagles Medium (Gibco, USA) supplemented with 10% heat-inactivated foetal bovine serum (Sigma-Aldrich, USA) at 37°C/5% CO_2_. Cells were placed into a 24-well plate at 1.5×10_5_ cells/well and allowed to adhere overnight. The following day, virus-containing samples (BALF, Lung and brain homogenates) were serially diluted in Modified Eagles Medium (MEM), cell culture supernatants removed from the VeroE6 cells and 250 µL of virus-containing samples was added to cell monolayers. After 1 hr, 250 *μ*L of 0.6% agar/MEM solution was gently overlaid onto samples and placed back into the incubator. At 72 hrs post-infection, each well was fixed with an equal volume of 8% paraformaldehyde solution (4% final solution) for 30 min at RT, followed by several washes with PBS and incubation with 0.025% crystal violet solution for 5 min at RT to reveal viral plaques.

### Hamster SARS-CoV-2 challenge

The hamster challenge studies were performed at the Vaccine and Infectious Disease Organization (VIDO, University of Saskatchewan, Canada). Male Golden Syrian hamsters (7-8 weeks old) were purchased from Charles River Laboratories (Charles River, Kingston, NY, U.S.A.). Hamsters were immunised i.m. twice three weeks apart in the quadriceps once with a total volume of 100 *μ*l. Each vaccine contained 5 or 20 *μ*g of chimeric spike antigen in endotoxin free PBS with SWE, or 100 *μ*l of Novavax NVX-CoV2373. Three weeks after the second immunisation, animals were challenged intranasally with 1×10_5_ TCID_50_ of SARS-CoV-2 variant Omicron (XBB.1.5) or 1×10_5_ TCID_50_ of SARS-CoV-2 variant Delta (B.1.617.2). Administration of the challenge virus was in both nares with 50 *μ*L/nare. Body weight of each animal was measured daily. Animals were euthanised either at 3 days post-challenge or at 10 days post-challenge. At necropsy, blood, lung tissues and nasal turbinate were collected for assessment of lesions, infectious virus quantification and histopathological examination. Infectious virus was determined by TCID_50_ analysis. Assays were conducted in 96-well plates using Vero 76 cells (ATCC CRL-1587). TCID_50_ was determined by microscopic observation of the cytopathic effect (CPE) on cells.

### Statistical analysis

The significance of differences between experimental groups was evaluated by either one-way ort two-way analysis of variance (ANOVA), with pairwise comparison of multi-grouped data sets achieved using Šidák’s, Tukey’s, Dunnett’s or Šidák’s *post-hoc* test, as indicated in the relevant figure legend. Where required, log transformation was used to obtain normal distribution and variance homogeneity prior to analysing data. Differences were considered statistically significant when p ≤ 0.05.

## Acknowledgements

This work was supported by the Coalition for Epidemic Preparedness Innovations (CEPI) as part of the Broadly Protective Beta-Coronavirus Program. We are grateful to the University of Sydney Drug Discovery Initiative and the Sydney Infectious Diseases Institute for support through their seed funding programs. We thank the Vaccine and Infectious Disease Organization for undertaking hamster challenge studies and helpful discussions; Charles Baily, Centenary Institute, Sydney, Australia for provision of lentivirus packaging and helper plasmids; Stuart Turville, Kirby Institute UNSW, Sydney, Australia for providing SARS-CoV-2 spike plasmid (Ancestral) and ACE2 293T cells; Nathaniel Landau, NYU Grossman School of Medicine, NY, USA for providing SARS-CoV-2 Delta spike plasmid. We thank Céline Lemoine, Falko Apel and Morgane Joessel from the Vaccine Formulation Institute, Switzerland for helpful discussions. We acknowledge the support of the University of Sydney Advanced Cytometry Facility and the animal facility at the Centenary Institute. Images created with Biorender.com where relevant.

## Author contributions

C.C, P.P. V.K.M, M.J.W, F.M.W and J.A.T conceived the study. C.C., M.D.J., E.C., C.A., E.E., J.T., S.A., S.M., N.G.H. Performed data acquisition. All authors undertook data analysis and interpretation of data. C.C, P.P. V.K.M, M.J.W, M.S., T.C., P.M.D, N.C, P.M.H, F.M.W and J.A.T contributed to resources and funding acquisition. The initial manuscript draft was prepared by J.A.T. and M.S. with input from all other authors. All authors approved the final version of the manuscript.

## References

1 Brice, Y., Morgan, L., Kirmani, M., Kirmani, M. & Udeh, M. C. COVID-19 Vaccine Evolution and Beyond. Neurosci Insights 18, 26331055231180543 (2023).

2 Mahrokhian, S. H., Tostanoski, L. H., Vidal, S. J. & Barouch, D. H. COVID-19 vaccines: Immune correlates and clinical outcomes. Hum Vaccin Immunother 20, 2324549 (2024).

3 Triccas, J. A. & Steain, M. C. Australia’s COVID-19 vaccine journey: progress and future perspectives. Microbiology Australia 45, 27–31 (2024).

4 Menegale, F. et al. Evaluation of Waning of SARS-CoV-2 Vaccine–Induced Immunity: A Systematic Review and Meta-analysis. JAMA Network Open 6, e2310650–e2310650 (2023).

5 Triccas, J. A., Kint, J. & Wurm, F. M. Affordable SARS-CoV-2 protein vaccines for the pandemic endgame. npj Vaccines 7, 89 (2022).

6 Temmam, S. et al. Bat coronaviruses related to SARS-CoV-2 and infectious for human cells. Nature 604, 330–336 (2022).

7 He, W. T. et al. Virome characterization of game animals in China reveals a spectrum of emerging pathogens. Cell 185, 1117–1129 e1118 (2022).

8 Marani, M., Katul, G. G., Pan, W. K. & Parolari, A. J. Intensity and frequency of extreme novel epidemics. Proc Natl Acad Sci U S A 118 (2021).

9 Hsieh, C. L. et al. Structure-based design of prefusion-stabilized SARS-CoV-2 spikes. Science 369, 1501–1505 (2020).

10 Calvaresi, V. et al. Structural dynamics in the evolution of SARS-CoV-2 spike glycoprotein. Nat Commun 14, 1421 (2023).

11 Khoury, D. S. et al. Neutralizing antibody levels are highly predictive of immune protection from symptomatic SARS-CoV-2 infection. Nat Med 27, 1205–1211 (2021).

12 Cromer, D. et al. Predicting vaccine effectiveness against severe COVID-19 over time and against variants: a meta-analysis. Nat Commun 14, 1633 (2023).

13 Cromer, D. et al. Neutralising antibody titres as predictors of protection against SARS-CoV-2 variants and the impact of boosting: a meta-analysis. Lancet Microbe 3, e52–e61 (2022).

14 Seifert, S. N. et al. An ACE2-dependent Sarbecovirus in Russian bats is resistant to SARS-CoV-2 vaccines. PLoS Pathog 18, e1010828 (2022).

15 Nie, J. et al. Functional comparison of SARS-CoV-2 with closely related pangolin and bat coronaviruses. Cell Discov 7, 21 (2021).

16 Cankat, S., Demael, M. U. & Swadling, L. In search of a pan-coronavirus vaccine: next-generation vaccine design and immune mechanisms. Cell Mol Immunol 21, 103–118 (2024).

17 Sette, A., Sidney, J. & Crotty, S. T Cell Responses to SARS-CoV-2. Annu Rev Immunol 41, 343–373 (2023).

18 Pino, P. et al. Trimeric SARS-CoV-2 Spike Proteins Produced from CHO Cells in Bioreactors Are High-Quality Antigens. Processes, 1539 (2020).

19 Counoupas, C. et al. High-Titer Neutralizing Antibodies against the SARS-CoV-2 Delta Variant Induced by Alhydroxyquim-II-Adjuvanted Trimeric Spike Antigens. Microbiol Spectr 10, e0169521 (2022).

20 Yao, Z., Zhang, L., Duan, Y., Tang, X. & Lu, J. Molecular insights into the adaptive evolution of SARS-CoV-2 spike protein. J Infect 88, 106121 (2024).

21 Zahradnik, J. et al. SARS-CoV-2 variant prediction and antiviral drug design are enabled by RBD in vitro evolution. Nat Microbiol 6, 1188–1198 (2021).

22 Aguilar-Bretones, M., Fouchier, R. A., Koopmans, M. P. & van Nierop, G. P. Impact of antigenic evolution and original antigenic sin on SARS-CoV-2 immunity. J Clin Invest 133 (2023).

23 Carabelli, A. M. et al. SARS-CoV-2 variant biology: immune escape, transmission and fitness. Nat Rev Microbiol 21, 162–177 (2023).

24 Focosi, D., Quiroga, R., McConnell, S., Johnson, M. C. & Casadevall, A. Convergent Evolution in SARS-CoV-2 Spike Creates a Variant Soup from Which New COVID-19 Waves Emerge. Int J Mol Sci 24 (2023).

25 Yang, S. et al. Fast evolution of SARS-CoV-2 BA.2.86 to JN.1 under heavy immune pressure. Lancet Infect Dis 24, e70–e72 (2024).

26 O’Hagan, D. T., van der Most, R., Lodaya, R. N., Coccia, M. & Lofano, G. “World in motion” - emulsion adjuvants rising to meet the pandemic challenges. NPJ Vaccines 6, 158 (2021).

27 Halfmann, P. J. et al. Broad protection against clade 1 sarbecoviruses after a single immunization with cocktail spike-protein-nanoparticle vaccine. Nat Commun 15, 1284 (2024).

28 Cohen, A. A. et al. Mosaic RBD nanoparticles protect against challenge by diverse sarbecoviruses in animal models. Science 377, eabq0839 (2022).

29 Evans, J. P. & Liu, S. L. Challenges and Prospects in Developing Future SARS-CoV-2 Vaccines: Overcoming Original Antigenic Sin and Inducing Broadly Neutralizing Antibodies. J Immunol 211, 1459–1467 (2023).

30 Schiepers, A. et al. Molecular fate-mapping of serum antibody responses to repeat immunization. Nature 615, 482–489 (2023).

31 Garg, R. et al. Efficacy of a stable broadly protective subunit vaccine platform against SARS-CoV-2 variants of concern. Vaccine (2024).

32 Kim, D. E., Chivian, D. & Baker, D. Protein structure prediction and analysis using the Robetta server. Nucleic Acids Res 32, W526–531 (2004).

33 Waterhouse, A. et al. SWISS-MODEL: homology modelling of protein structures and complexes. Nucleic Acids Res 46, W296–W303 (2018).

34 Efimov, V. P. et al. Fibritin encoded by bacteriophage T4 gene wac has a parallel triple-stranded alpha-helical coiled-coil structure. J Mol Biol 242, 470–486 (1994).

35 Jin, J. et al. Accelerating the clinical development of protein-based vaccines for malaria by efficient purification using a four amino acid C-terminal ‘C-tag’. Int J Parasitol 47, 435–446 (2017).

36 Counoupas, C. et al. A single dose, BCG-adjuvanted COVID-19 vaccine provides sterilising immunity against SARS-CoV-2 infection. NPJ Vaccines 6, 143 (2021).

37 Counoupas, C. et al. Mycobacterium tuberculosis components expressed during chronic infection of the lung contribute to long-term control of pulmonary tuberculosis in mice. NPJ Vaccines 1, 16012 (2016).

