## Supporting information for "A Single Chimeric Spike Antigen Induces Pan-Sarbecovirus Immunity"

### Counoupas *et. al.*

**Supplementary Table 1:** Characteristics of the CSA05 antigen (1g/L)

| Category | Characteristic | Storage Temperature | (T-1) Δt = -1 day | | (T0) Δt = 0 | | (T1) Δt = 4 weeks | | (T2) Δt = 8 weeks | | (T3) Δt = 16 weeks | | (T4) Δt = 25 weeks | | Unit |
|---|---|---|---|---|---|---|---|---|---|---|---|---|---|---|---|
| | | | AV | SD | AV | SD | AV | SD | AV | SD | AV | SD | AV | SD | |
| Quantification | Protein concentration (A280) | **4°C** | 989.03 | 9.27 | 997.55 | 13.33 | 997.27 | 4.28 | 987.17 | 8.82 | 993.48 | 8.93 | 989.89 | 9.30 | mg/L |
| | | **-20 °C** | nd | nd | 1011.15 | 11.53 | 996.46 | 1.84 | 1000.83 | 6.91 | 979.54 | 38.53 | 1016.04 | 9.47 | mg/L |
| | | **-80 °C** | nd | nd | 1016.01 | 18.05 | 998.13 | 5.08 | 1006.47 | 3.89 | 1003.71 | 12.75 | 1004.04 | 12.64 | mg/L |
| Product-related impurities | Purity (SEC-UV) | **4°C** | 96.6% | 0.2% | 97.5% | 0.2% | 97.3% | 0.1% | 97.5% | 0.0% | 97.5% | 0.0% | 97.3% | 0.0% | % |
| | | **-20 °C** | nd | nd | 97.5% | 0.2% | 97.2% | 0.0% | 97.3% | 0.0% | 97.3% | 0.0% | 96.7% | 0.1% | % |
| | | **-80 °C** | nd | nd | 97.1% | 0.2% | 97.0% | 0.1% | 97.3% | 0.0% | 97.3% | 0.0% | 97.1% | 0.0% | % |
| Integrity | Size (SDS-PAGE) | **4°C** | 150.01 | 1.30 | 152.66 | 4.81 | 149.75 | 0.55 | 152.18 | 3.88 | 152.29 | 3.04 | 149.01 | 1.04 | kDa |
| | | **-20 °C** | nd | nd | 150.57 | 3.33 | 150.06 | 0.60 | 151.15 | 3.04 | 150.75 | 3.71 | 148.41 | 0.79 | kDa |
| | | **-80 °C** | nd | nd | 148.57 | 1.67 | 149.39 | 0.37 | 148.56 | 1.32 | 147.81 | 1.99 | 150.22 | 0.78 | kDa |
| Potency | Dissociation constant (BLI) | **4°C** | nd | nd | 2.76E-09 | 9.85E-10 | 2.75E-09 | 1.05E-09 | 2.89E-09 | 9.82E-10 | 4.47E-09 | 2.64E-09 | 3.19E-09 | 1.26E-09 | mol L-1 |
| | | **-20 °C** | nd | nd | 2.73E-09 | 1.12E-09 | 2.72E-09 | 9.81E-10 | 2.79E-09 | 1.00E-09 | 4.00E-09 | 1.46E-09 | 3.03E-09 | 1.16E-09 | mol L-1 |
| | | **-80 °C** | nd | nd | 2.61E-09 | 1.03E-09 | 2.58E-09 | 1.00E-09 | 2.69E-09 | 1.03E-09 | 4.16E-09 | 2.03E-09 | 3.08E-09 | 1.15E-09 | mol L-1 |

**Supplementary Table 2:** Individual neutralising antibody values (IU/ml) used for Figure 5 spider plots.

| | | pre-boost | | | | | | post-boost | | | | | |
|---|---|---|---|---|---|---|---|---|---|---|---|---|---|
| **Ancestral** | CoVEXS5 (1 µg) | 96876.32 | 27299.48 | 18163.39 | 9802.84 | 10523.38 | 20858.4 | 455432.8 | 455432.8 | 455432.8 | 32066.45 | 46767.65 | 109572.5 |
| | CoVEXS5 (5 µg) | 29508.92 | 15422.04 | 26771.9 | 20084.77 | 12022.8 | 28271.4 | 455432.8 | 198585.1 | 37925.76 | 58449.01 | 62781.36 | 53706.88 |
| | NVX | 94225.48 | 40681.82 | 21967.69 | 16266.81 | 16504.48 | 31424.03 | 111561.2 | 455432.8 | 113135.1 | 79503.76 | 44565.82 | 455432.8 |
| **Delta** | CoVEXS5 (1 µg) | 89351.86 | 18513.65 | 12510.54 | 9044.215 | 7175.345 | 16491.11 | 455432.8 | 455432.8 | 72176.21 | 20692.46 | 31261.13 | 74975.09 |
| | CoVEXS5 (5 µg) | 22316.35 | 12348.24 | 24038.17 | 16797.45 | 5499.76 | 15281.63 | 455432.8 | 455432.8 | 43113.19 | 48033.65 | 50934.37 | 44436.99 |
| | NVX | 31391.2 | 18822.3 | 14073.91 | 11722.73 | 9884.858 | 11659.57 | 42598.42 | 57625.67 | 90265.12 | 42825.02 | 44150.5 | 128003.6 |
| BA.1 | CoVEXS5 (1 µg) | 2824.661 | 3579.248 | 379.1398 | 2750.147 | 2496.094 | 4205.841 | 18454.7 | 30043.96 | 14142.88 | 15063.72 | 11310.08 | 26919.62 |
| | CoVEXS5 (5 µg) | 13802.84 | 3211.729 | 5407.28 | 2458.78 | 3159.959 | 3248.221 | 249266.8 | 215401.4 | 49261.27 | 32061.92 | 27184.67 | 70696.24 |
| | NVX | 909.4028 | 1145.749 | 5124.777 | 2436.94 | 1803.657 | 1855.32 | 51308.27 | 6128.62 | 107836.9 | 18466.25 | 57144.33 | 35959.71 |
| **BA.2** | CoVEXS5 (1 µg) | 2308.145 | 3539.243 | 1657.608 | 3045.742 | 1925.983 | 550.0475 | 17004.04 | 42735.45 | 46157.83 | 39746.84 | 46444.95 | 32331.67 |
| | CoVEXS5 (5 µg) | 4766.009 | 1326.655 | 2071.357 | 3169.475 | 1823.231 | 5027.635 | 61910.17 | 122724.6 | 29509.48 | 62877.56 | 51037.38 | 62791.77 |
| | NVX | 184.0406 | 1365.266 | 11703.02 | 4077.107 | 1838.433 | 823.6102 | 7968.854 | 3970.004 | 125028.3 | 15410.17 | 72519.94 | 15447.61 |
| **BA.5** | CoVEXS5 (1 µg) | 1068.437 | 7190.204 | 738.6908 | 4848.538 | 2602.565 | 824.1129 | 17495.4 | 35098.66 | 47185.94 | 20016.41 | 40213.75 | 29404.38 |
| | CoVEXS5 (5 µg) | 6585.735 | 2777.418 | 5428.453 | 6780.419 | 1911.732 | 3783.096 | 75441.12 | 113727 | 32211.65 | 35133.18 | 39906.15 | 44956.71 |
| | NVX | | 786.4258 | 7985.578 | 2880.266 | 1629.14 | | | 3435.377 | 114303.3 | 12153.9 | 65359.46 | |
| **BQ.1.1** | CoVEXS5 (1 µg) | 30469.1 | 3661.044 | 1215.864 | 1570.134 | 5813.534 | 8599.605 | 48766.17 | 14433.95 | 8521.645 | 3029.531 | 13248.04 | 39230.22 |
| | CoVEXS5 (5 µg) | 23065.75 | 4721.265 | 2076.561 | 2460.31 | 3991.835 | 7896.165 | 184692.6 | 19894.21 | 3307.216 | 4614.607 | 15363.11 | 28092.64 |
| | NVX | 3117.002 | 2475.432 | 2980.188 | 2385.194 | 1816.025 | 2712.822 | 3494.38 | 11141.21 | 13752.76 | 5719.467 | 7410.659 | 8257.617 |
| **XBB.1.5** | CoVEXS5 (1 µg) | 2240.4707 | 208.24545 | 1524.6236 | 641.28954 | 208.24545 | 1119.6723 | 13531.376 | 13822.029 | 9500.0807 | 1629.7806 | 1116.9416 | 26093.536 |
| | CoVEXS5 (5 µg) | 208.24545 | 208.24545 | 208.24545 | 208.24545 | 208.24545 | 208.24545 | 5254.6428 | 2835.3638 | 3427.1646 | 624.73634 | 2686.4544 | 1285.0342 |
| | NVX | 2497.6479 | 3372.2302 | 4404.8695 | 2158.5933 | 4000.4489 | 1274.0961 | 14609.704 | 10505.139 | 5449.1179 | 4463.937 | 2775.7336 | 5999.347 |
| **JN.1** | CoVEXS5 (1 µg) | 80 | 80 | 80 | 80 | 80 | 129.7123 | 192.1634 | 101.6503 | 140.6708 | 80 | 782.9186 | 80 |
| | CoVEXS5 (5 µg) | 80 | 80 | 80 | 80 | 80 | 80 | 698.0179 | 241.8437 | 355.6101 | 80 | | 493.4394 |
| | NVX | 80 | 80 | 101.2538 | 80 | 80 | 557.0684 | 80 | 222.3235 | 367.1661 | 91.46757 | 414.6681 | 188.0947 |
| **BtKY72** | CoVEXS5 (1 µg) | 4006.099 | 698.2943 | 929.6379 | 2996.239 | 3576.672 | 1029.277 | 27255.49 | 14520.1 | 27126.34 | 22847.09 | 27659.02 | 18977.89 |
| | CoVEXS5 (5 µg) | 1430.208 | 446.4887 | 1367.048 | 737.7402 | 3288.367 | 16370.93 | 31565.88 | 36247.02 | 12894.74 | 33507.45 | | 59698.64 |
| | NVX | 4142.932 | 7369.692 | 1517.536 | 905.9774 | 5610.968 | 4079.575 | 3461.886 | 27296.84 | 23938.83 | 26187.28 | 94557.88 | 46877.7 |
| **Pangolin GX** | CoVEXS5 (1 µg) | 210.7543 | 208.2454 | 208.2454 | 208.2454 | 208.2454 | 384.5675 | 937.9852 | 557.6339 | 520.973 | 243.2911 | 575.4212 | 1584.829 |
| | CoVEXS5 (5 µg) | 407.4198 | 208.2454 | 272.1946 | 208.2454 | 391.1112 | 460.7668 | 1676.128 | 1099.243 | 372.5744 | 1111.85 | | 1910.198 |
| | NVX | 202.693 | 451.9144 | 434.1275 | 596.0136 | 425.4778 | 456.7981 | 449.6861 | 674.3939 | 1106.64 | 1078.055 | 1631.586 | 1151.348 |
| **Pangolin GD** | CoVEXS5 (1 µg) | 174960 | 29103.15 | 21527.08 | 27702.39 | 53757.31 | 8862.575 | 174960 | 174960 | 174960 | 174960 | 174960 | 174960 |
| | CoVEXS5 (5 µg) | 98495.47 | 9511.927 | 37134.58 | 15524.69 | 9760.342 | 92825.12 | 174960 | 174960 | 174960 | 174960 | 174960 | 174960 |
| | NVX | 58320 | 143737.3 | 150458.8 | 26888.47 | 112061.4 | 66264.86 | 238211.7 | 193063.7 | 174960 | 174960 | 174960 | 174960 |
| **SARS-CoV-1** | CoVEXS5 (1 µg) | 19440 | 1166.39 | 579.5653 | 701.8059 | 666.0416 | 917.3573 | 58320 | 4821.134 | 22675.06 | 1583.346 | 4185.962 | 861.3168 |
| | CoVEXS5 (5 µg) | 747.6452 | 712.7206 | 672.5691 | 579.3301 | 600.0177 | 357.1256 | 24747.41 | 2034.647 | 1752.596 | 1600.738 | 600 | 685.3649 |
| | NVX | 502.7475 | 1400.645 | 815.1846 | 982.0348 | 1512.292 | 1520.511 | 914.4383 | 2368.842 | 9957.547 | 3004.114 | 2452.374 | 16382.11 |
| **WIV-1** | CoVEXS5 (1 µg) | 8704.331 | 2166.059 | 1437.903 | 2213.198 | 1631.148 | 1783.995 | 174960 | 9051.696 | 11647.22 | 174960 | 15341.49 | 14835.51 |
| | CoVEXS5 (5 µg) | 1690.569 | 1422.634 | 1816.432 | 1020.925 | 1627.842 | 1900.705 | 91739.13 | 9520.653 | 2139.719 | 6859.563 | 1627 | 3736.799 |
| | NVX | 6411.75 | 3860.673 | 5039.667 | 1951.513 | 3743.774 | 1828.118 | 41391.21 | 10968.69 | 6427.545 | 27824 | 8080.124 | 7160.002 |
